# Engineering Nanowires in Bacteria to Elucidate Electron Transport Structural-Functional Relationships

**DOI:** 10.1101/2022.10.06.510814

**Authors:** Ben Myers, Francesco Catrambone, Stephanie Allen, Phil J Hill, Katalin Kovacs, Frankie J Rawson

**Affiliations:** Bioelectronics Laboratory, Regenerative Medicine and Cellular Therapies, School of Pharmacy, Biodiscovery Institute, The University of Nottingham, Nottingham, NG7 2RD, UK; BBSRC/EPSRC Synthetic Biology Research Centre, School of Life Sciences, Biodiscovery Institute, University of Nottingham, NG7 2RD, United Kingdom; Molecular Therapeutics and Formulation Division, School of Pharmacy, University Park, The University of Nottingham, Nottingham, NG7 2RD, UK; Division of School of Microbiology, Brewing and Biotechnology, School of Biosciences, Sutton Bonington Campus, University of Nottingham, LE12 5RD, United Kingdom

## Abstract

Bacterial pilin nanowires are protein complexes, suggested to possess electroactive capabilities forming part of the cells’ bioenergetic programming. Their role is thought to be linked to facilitating electron transfer with the external environment to permit metabolism and cell-to-cell communication. There is a significant debate, with varying hypotheses as to the nature of the proteins currently lying between type-IV pilin-based nanowires and polymerised cytochrome-based filaments. Importantly, to date, there is a very limited structure-function analysis of these structures within whole bacteria. In this work, we engineered *Cupriavidus necator* H16, a model autotrophic organism to express differing aromatic modifications of type-IV pilus proteins to establish structure-function relationships on conductivity and the effects this has on pili structure. This was achieved *via* a combination of high-resolution PeakForce tunnelling atomic force microscopy (PeakForce TUNA™) technology, alongside conventional electrochemical approaches enabling the elucidation of conductive nanowires emanating from whole bacterial cells for the first time. This work is the first example of functional type-IV pili protein nanowires produced under aerobic conditions using a *CN* chassis. This work has far-reaching consequences in understanding the basis of bio-electrical communication between cells and with their external environment.

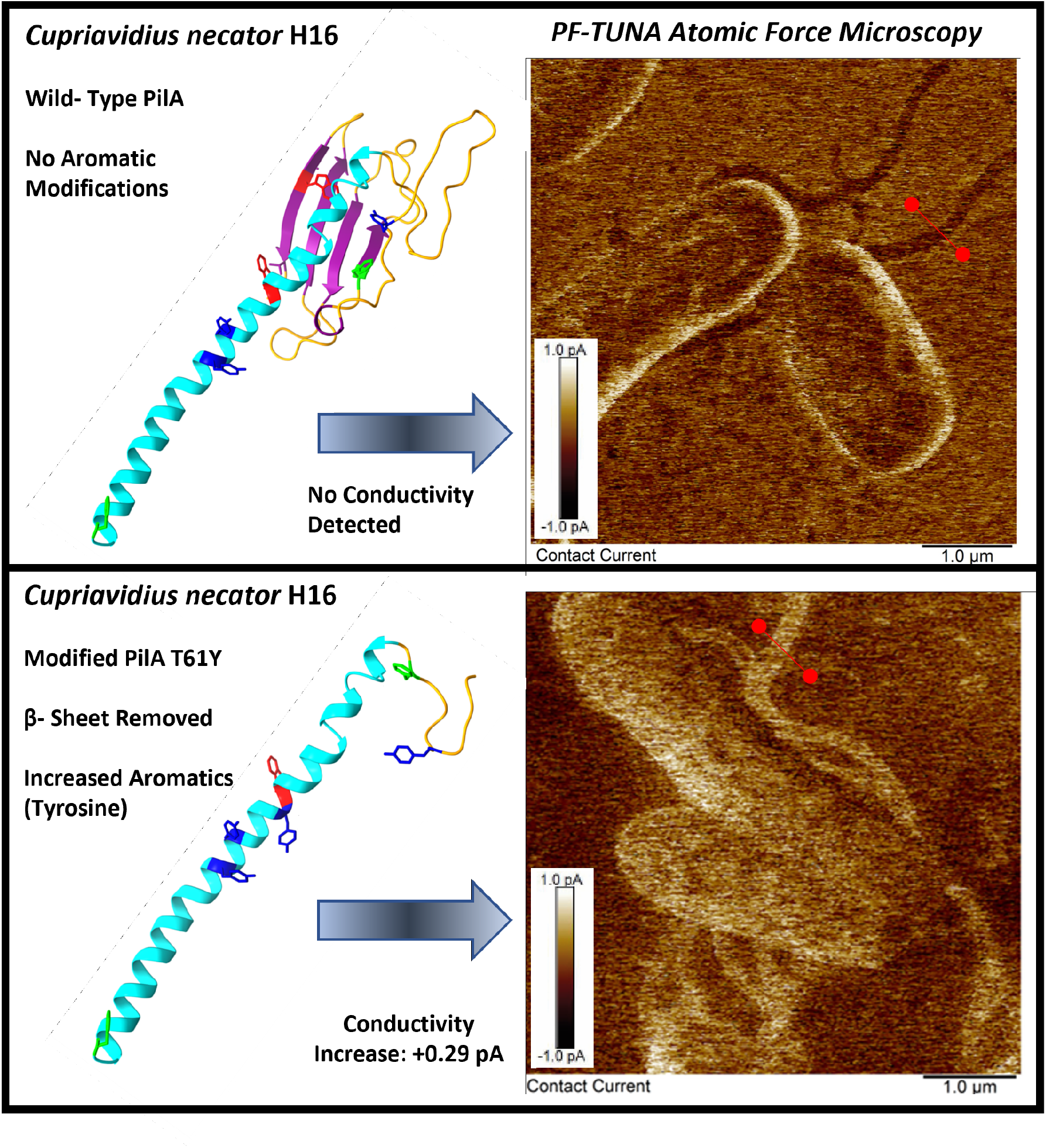

Graphical abstract displaying theoretical PilA monomer models (left), PeakForce TUNA atomic force microscopy contact current images (right) of wild-type (top) and modified with increased tyrosine content (bottom) PilA filaments expressed by *Cupriavidus necator* H16 cells.

## Introduction

The discovery and study of bacterial protein nanowires has revolutionised our understanding of microbial electrochemical communication mechanisms. Conductive nanowires were first observed in the Gram-negative *Geobacter sulfurreducens* (*GS*) ^1^ and have been identified across several phyla ^2^. Microbial nanowires are thought to be part of the mechanism by which electrons are transported outside the cell, connecting their internal redox centres with external reductive or oxidative sources. In nature, these structures are used by heterogeneous and pure microbial communities to extend the distance of electrochemical interactions between cell-to-mineral and cell-to-cell connections^3^.

Their electrical properties have suggested uses as potential biological electronic material ^4^, based on conductivity comparable to synthetic metallic structures ^5^, non-toxicity, and ability to maintain these characteristics across varying environmental conditions ^6^. Moreover, the expression of conductive nanowires in heterologous hosts may provide opportunities in producing novel exoelectrogenic microbial chassis for bioelectrochemical systems applications ^7,8^.

However, the mechanism of electron transport and the identity of the proteins involved are less clearly understood. The term ‘Microbial Nanowires’ refers to conductive, extracellularly-projected filamentous protein assemblies produced by microorganisms to electrochemically communicate with their environment. These structures are thought to be comprised of either repeating subunits of a type-IV pilin monomer ^9^, or polymerised cytochromes containing a stacked inner core of heme-ring structures ^10^. Compelling evidence of both molecular identities has been demonstrated ^9–14^, and their specific protein composition has been hotly contested ^15,16^. Indeed, it is feasible that *GS* produces multiple forms of extracellularly projected conductive proteins, in addition to the array of soluble mediator compounds and surface-bound redox-active proteins also shown to exist ^17,18^.

Nevertheless, extensive experimental evidence supports the type-IV pilin mechanistic model of conductivity ^5,8,14,19–21^, where nanowires are constituted of polymerised PilA subunits with aromatic amino acids located at specific positions, bringing overlapping π-π orbitals of neighbouring aromatic rings within the spatial distance to allow electron flux ^22^. Replacing aromatic amino acids for the non-aromatic alanine produced the *GS* Aro-5 strain, with a diminished Fe (ii) oxide reduction ability ^9^, alongside reduced pilus conductivity measured via atomic force microscopy (AFM) based current-voltage spectroscopy. In contrast, replacing native residues for tryptophan, which possesses a double aromatic ring, increased pilus conductivity by over 500% ^5^. Type-IV pilin nanowires capable of electron transport have also been expressed in a range of heterologous bacterial hosts, both by expressing the entire *GS* PilA monomer sequence ^6^ or modifying the aromatic content of native, nonconductive type-IV pili ^21^. So far, recombinant conductive nanowires have been achieved in *GS* ^5^, *Pseudomonas aeruginosa (PA)* ^21^, *Shewanella Oneidensis* ^23^ and *Escherichia coli* ^6^, all Gram-negative bacteria possessing type-IV pili generally used for surface adherence and twitching motility ^24^.

Providing further evidence of the relationship between aromaticity and pilus conductivity, a 2022 study also demonstrated the first example of a conductive protein based on the type-I pilus architecture of *E. coli* ^14^, via aromatic modification. These findings are difficult to align with the observation that the conductive filaments originally identified as pili were comprised of polymerised cytochromes ^15^.

Conductive AFM (C-AFM) is the most frequently cited method for measuring the electrical properties of individual pili: with a range of methods to locate and image conductive filaments, including using a pilus ^25^ or biofilm ^23^ to bridge the gap between two electrodes, detached and purified pili deposited on a substratum ^6^ and whole-cell AFM analysis of filaments directly protruding from cells ^11^.

However, across these methodologies, no high-resolution images of the conductive data channel have been published to date. Instead, current-voltage spectroscopy is typically used to determine the conductivity and resistance of specific regions along the pili following the acquisition of an image of the sample topography. While these methods are useful to extract quantitative data on conductivity, it is debated whether extensive processing involved in sample preparation methods can cause the artificial polymerisation of individual monomers during precipitation and reconstitution, creating artefacts and synthetically modifying observed conductivities ^16^.

In contrast to the extensive reporting of current-voltage curves in the literature, to our knowledge, only a handful of studies have released C-AFM current maps of type-IV pili ^1,21,26^ and to date, current data images of filaments emerging from intact cells have not been published. The first current data images of sheared pili deposited on a substratum were released in 2005 ^1^, and other researchers have expanded this work via the visualisation of precipitated pili ^21^ and drop-cast bacterial cultures ^26,27^, although still without the wide-scale current data maps of intact conductive filaments directly emerging from cells. Such data is vital to address concerns regarding how the preparation of microbial nanowires affects their conductivity ^15,28,29^ and has been achieved in other organisms and proteins. For example, high-resolution C-AFM images of conductive cable bacteria ^30^ and hippocampal neurons ^31^ have been achieved, validating this technique as suitable to produce wide-scale conductive mapping of whole cells.

The reason for the lack of this type of data for conductive microbial nanowires, either pili or cytochrome-based, is unexplained, but likely attributable to technical difficulties in maintaining a balance between the spatial positioning of the AFM cantilever to capture electrical properties, while minimising sample damage or tip contamination. This is challenging to achieve using traditional contact-mode C-AFM, as the high lateral forces applied by dragging the cantilever across the surface frequently destroy biological samples. The use of PeakForce tunnelling atomic force microscopy (PeakForce TUNA™) overcomes this limitation by gently oscillating the cantilever (approx. 1kHz, 200nm amplitude) so that the tip intermittently contacts the sample surface during imaging ^32^. This decreases the contact time between cantilever and sample, reducing damaging lateral forces and limiting adhesion of cellular material to the cantilever tip whilst allowing simultaneous measurement of topographic and electrical data within one field of view.

This study aims to address in more detail the hypothesis that type-IV pilus aromaticity contributes to filament conductivity. We do this by elucidating the structure-function relationship of pili, resolving the first high-resolution PeakForce TUNA AFM current data and correlating the electrochemical behaviour of conductive type-IV filaments in *Cupriavidus necator* H16 *(CN)*, an industrially relevant facultative heterotroph ^33^, capable of oxidising a wide range of nutrient and energy sources, including CO_2_ and H_2_ as the sole source of carbon and energy. Due to its metabolic versatility and the availability of genetic tools ^34^, this organism has been successfully engineered to produce several industrially relevant molecules, such as solvents, fuels, terpenes and biopolymers ^35^, which makes this organism a promising candidate for microbial electrosynthesis using bioelectrochemical systems (BES), where the energy required for microbial growth and product synthesis can be derived from renewable electricity. We expressed conductive pili in *CN* via modification of its native type-IV pilus sequence, increasing its aromatic content. By showing these filaments directly emerging from intact bacterial cells, we identify structure-function relationships of the pili and demonstrate how the electron transfer efficiency can be modulated. This has implications for developing new bioelectronic materials and incorporating such bacteria into bioelectronics systems such as microbial fuel cells

In terms of the suitability of *CN* as a conductive pili expression host, it expresses a native type-IV PilA monomer structurally related to *GS* PilA, lacking aromatic amino acids in four key locations throughout the N-terminal α-helix. Our hypothesis predicts *CN*’s wild-type PilA to be non-conductive, given its aromatic content and distribution. Therefore, if pilus conductivity is attributable to aromatic content, replacing the native residues for aromatics at the same key locations present on the *GS* monomer should lead to an increase in *CN* PilA filament conductivity. Moreover, removing β-sheet regions to produce a PilA monomer comprised of only α-helix, is theorised to result in marginal conductivity increase by bringing aromatic residues to closer spatial distances when assembled into the major pilus ultrastructure.

## Results

### Design of native *CN* PilA monomers with variable aromatics content and localisation

To establish the mechanism affecting pili conductivity, we initially sought to engineer *CN* native type-IV PilA monomers to be expressed as variants with an array of aromatic amino acid substitutions as seen in Figure 1. *CN* PilA (Figure 1A) is structurally like that of *PA* (Figure 1B), *GS* (Figure 1C) and *GM* (Figure 1D) but with a difference in chain length (*CN*= 161 AA, *PA*= 143 AA, *GS*= 61AA, *GM*= 59AA).

**Figure 1.**
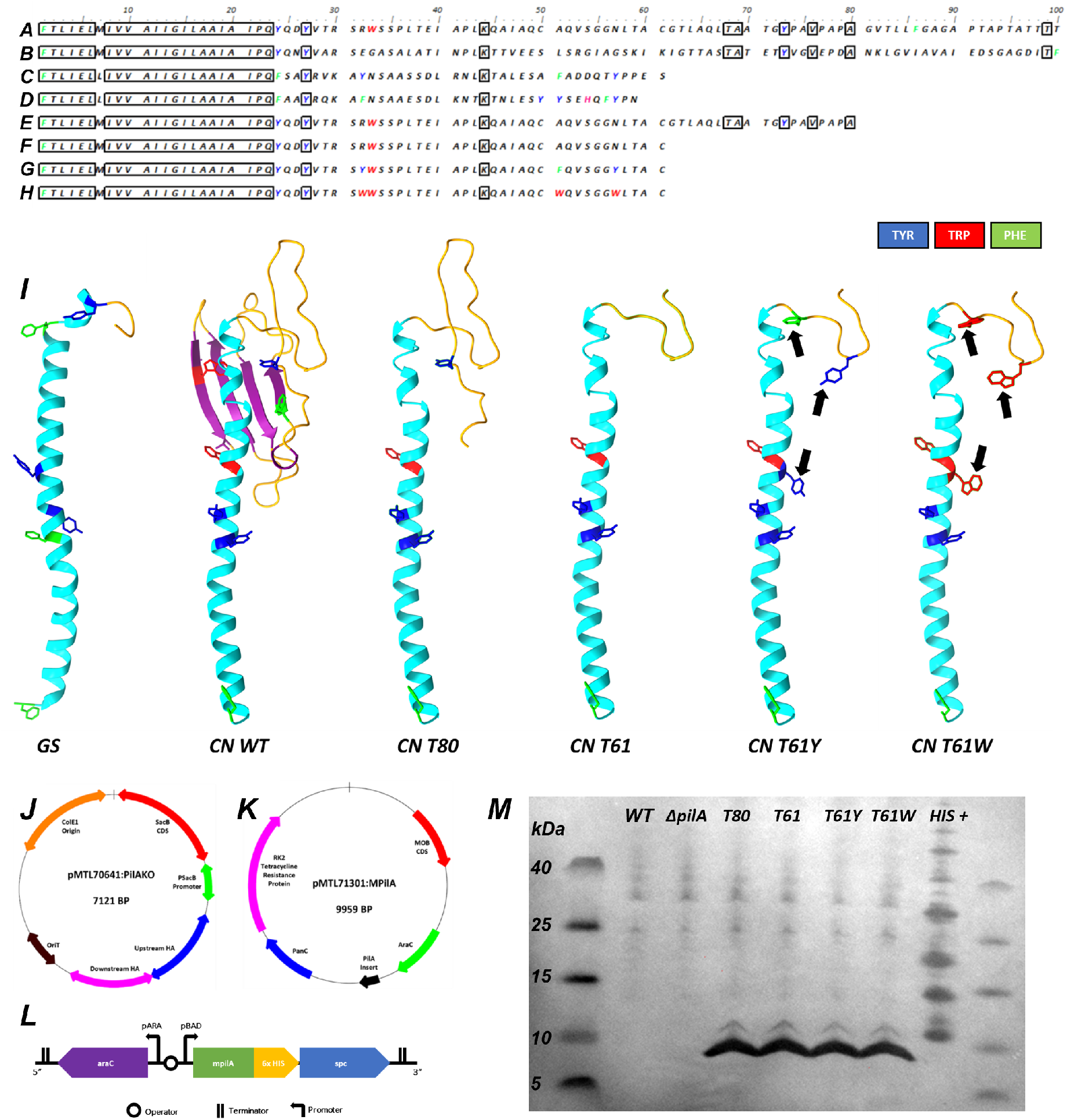
Design, expression, and verification of modified PilA (mPilA) monomer sequences in *Cupriavidus necator* H16. A-D) Native PilA monomer sequence alignment of A) *Cupriavidus necator* H16 *(CN)*; B) *Pseudomonas aeruginosa* (*PA*); and the conductive monomers of C) *Geobacter sulfurreducens* (*GS*) and D) *Geobacter metallidurans* (*GM*). E-H) Modified *CN* PilA sequences, produced in this study: E) *CN:* T80, and F) *CN*: T61, PilA monomers with structural but without aromatic modification. G) *CN*: T61Y, and H) *CN*: T61W monomer sequences, with both structural modifications and increased aromatic amino acid content. Sequences A and B (*CN*, PA) are limited to 100 characters for clarity (from 147 and 143, respectively). Boxed residues indicate homologous regions and coloured residues indicate aromaticity. Alignment produced via Bio Edit. I) Predictive models of type-IV PilA monomer structures. Ii) *GS* WT; Iii) *CN* WT; Iiii) *CN*: T61, with C-terminal β-sheet removed but lacking aromatic modification; Iiii) *CN*: T61W, an adaptation of T61 with native amino acids replaced for tryptophan at locations 32, 51, 57. Aromatic residues are highlighted as red (tryptophan), blue (tyrosine) or green (phenylalanine) branches, and black arrows indicate locations of amino acid mutation. Models produced via Chimera. J) pMTL70641: ΔpilA knockout vector; K) Modular pMTL71301: mPilA expression vector, with mPilA expression operon shown as a black arrow at 4.5 KB. Plasmid maps produced via ChemBioDraw. L) Expanded diagram of the mPilA expression operon, with variable mPilA insert gene (green) for each variant, preceded by *CN*’s PilA recognition sequence and in-frame C-terminal 6xHIS fusion. For the non-expressing negative control strain, this region was emitted. M) SDS-PAGE-Western blot of sheared pili extract showing HIS-tagged protein for mPilA variants at ∼9KDa, which is not present in the WT or ΔpilA strains.

Their N-terminus includes a short Sec signal recognition sequence, and it is structured as an α-helix comprising approximately 60 AA (Figure 1li). The conductivity of *GS* and *GM* pili is thought to be attributable to the closer spatial positioning of aromatic residues along the monomer sequence ^19^ (Figure 1li). The *PA* and *CN* monomers possess additional β-sheet and random coil regions which are not present in their counterparts (Figure 1lii), and it is speculated that their influence on the nanowire ultrastructure may increase distances between aromatic residues, interfering with the electron flux. Previous research suggested that cleaving PilA monomers to include only α-helix domains can lead to an increase in conductivity ^21^.

For this reason, all the modified pilA sequences (mPilA, Figure 1E-H) were truncated in their C-terminal: the mpilA sequence designated T80 was truncated from 100 to 80 AA residues leaving a portion of C-terminal β-sheet, and the sequence designated T61 was further truncated to a 61 AA length, leaving only the α-helix. PilA variant T61-Y is a derivative of T61 with three amino acid substitutions at R32Y, A51F, N57Y, while T61-W variant has three amino acid substitutions at R32W, A51W and N57W. We theorised that variants T80 and T61 will display a marginal-to non-existent conductivity, while the increased aromatic content of T61-Y and T61-W variants may show increased electron flux.

### *CN* strain engineering, cloning and expression of modified PilA nanowires

A *CN* H16 chassis with a knock-out of the native type-IV pilus monomer gene was generated as a background strain to avoid cross-reactions between the native and modified variants with the endogenous type-IV pili machinery. For this reason, the *pilA* gene in locus A0550 was deleted *via* homologous recombination using the plasmid pMTL70641PilAKO (Figure 1J). The successful deletion was verified *via* whole-genome sequencing, producing the *CN* Δ*pilA* strain.

The modified *pilA* (*MpilA*) inserts were assembled onto the pMTL71301 modular vector ^34^, producing plasmids pMTL71301:mPilA:T80 to T61W (Figure 1K). Briefly, the *MpilA* ORFs consisted of (from 5’ to 3’) the *CN* native PilA Sec signal sequence fused in-frame with different *MpilA* inserts (T80, T61, T61-Y and T61-W) with an in-frame C-terminal 6xHIS fusion (Figure 1L) and the small protein chaperone (SPC) protein of *GS*, which has been found to improve pilA monomer stability and assist its transport to the T4P construction apparatus located at the cytoplasmic membrane ^36^.MPilA expression was controlled using the P_BAD_ arabinose-inducible promoter and AraC transcriptional regulator, together forming the araBAD inducible system. Finally, a pMTL71301 empty plasmid lacking the modified *pilA* cargo was used as a negative control (Figure 1K,L).

Plasmids were transformed into the *CN* Δ*pilA* chassis, and the expression was determined via western blot detection of sheared and purified pili, using an anti-6xHIS antibody (Figure 1M). Physical shearing of surface-expressed filaments from cells was performed via centrifugation and passing through a fine hypodermic needle, as previously described ^37^. Western blots of mPilA variants showed 6xHIS-tagged protein bands at ∼9KDa for all mPilA variants (Figure 1M) but were not detected in *CN* WT or CN Δ*pilA* control chassis expressing an empty pMTL71301 plasmid.

### PeakForce TUNA Atomic Force Microscopy Analysis of strains expressing modified type-IV pili

To investigate the effect of modifying the aromatic content of the *CN* PilA sequence, PeakForce TUNA AFM ^38^, a propriety method of tunnelling conductive atomic force microscopy was used. Figure 2 provides an overview of the PeakForce TUNA AFM data captured for all CN strains. A *CN* H16 WT strain, as well as *CN* Δ*pilA* were used as negative controls to determine the conductivity of native filaments. For each strain, the left image is the ‘PeakForce Error’ image, as this provides the clearest visual representation of the sample topography and as such was used to identify pili-rich locations within the sample. The right image provides the simultaneously captured contact current images for the same sample area, where brighter features reveal a higher recorded current. Topographical data, including that used to calculate pili dimensions, is displayed in Supplementary Information (SI) Figure 1.

**Figure 2.**
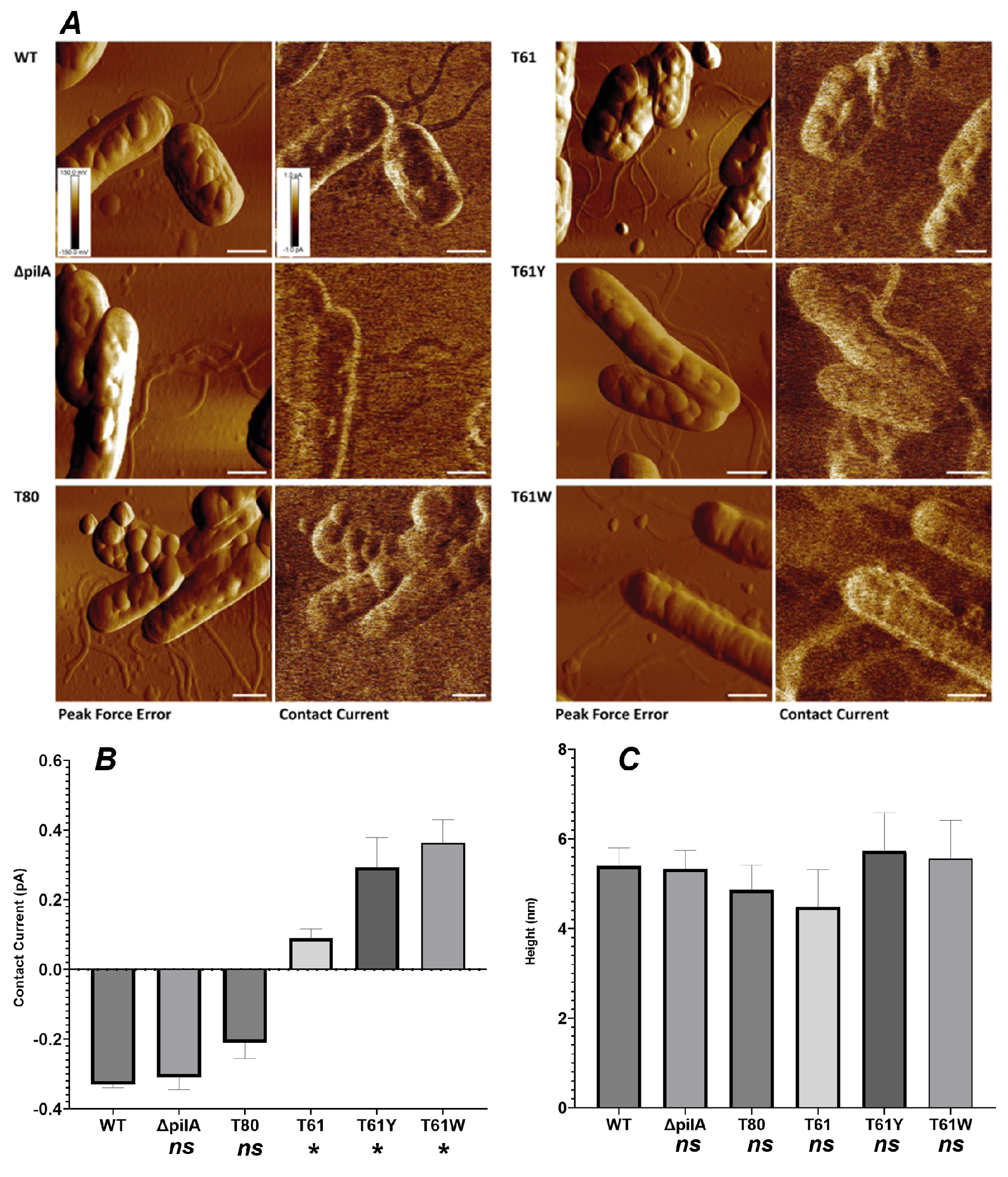
PeakForce TUNA AFM of *C. necator* H16 strains expressing wild-type (WT; *ΔpilA*) and modified (T80; T61; T61Y; T61W) type-IV pilA filaments. A: Peak Force Error, and Contact Current data images, with the corresponding strain labelled to the left of each image. Z-scale shown in the lower-left corner of WT, aligned across all images to a scale of +150 mV to -150 mV (Peak Force Error), and +1pA to -1pA (Contact Current). The white bar in the bottom right of each image represents a lateral scale of 1 µm. Charts b and c display Contact current (b), and Height (c) line profile gradient analysis. B: Contact current peak size, corresponding to pA increase or decrease over the baseline. C: Pili filament cross-sectional height. All variants had a mean filament morphology of 4.5-5.7 nm, and differences between WT and engineered pili height were not significant (p>0.05). For each data channel, nine measurements were taken in total for each variant, specifically three profiles drawn across three individual filaments (n=9 N=3). The mean of each filament is displayed in SI Figures 1 and 2. Line profile analysis was performed via Gwyddion39. Statistical analysis performed via one-way ANOVA of experimental sample against the WT (p<0.05), via Graphpad Prism. Significantly different comparisons are indicated by *, and *ns* indicates comparisons with p>0.05.

Surface-expressed filaments were observed in all *CN* strains, interestingly even in the *CN* Δ*pilA* strain with native pilA monomer deleted (Figure 2). This latter phenotype could indicate the presence of multiple genes encoding for unannotated type-IV pilus monomers, or other uncharacterised surface-expressed filaments such as a type-I pili or flagella. Nevertheless, no current increase over that of the background was observed in strains without the structural or aromatic modifications theorised to produce a conductive pilus, indicated by the darker colour of pili regions within the current data images compared to the substrate (Figure 2, WT; Δ*pilA*; T80). Contact current line profile analysis displayed mean decreases of -0.33 pA (SD ± 0.01) and -0.31 pA (SD ± 0.03), compared to the glass substratum, for the control strains *CN* H16 and *CN* Δ*pilA* respectively (Figure 2b). A decrease of -0.21pA (SD ± 0.05) was observed for *CN* T80 (Figure 2b). The T61 strain, expressing a pilus cleaved to only incorporate the α-helix region with no aromatic modifications, resulted in a contact current increment of 0.09pA (SD ± 0.03) (Figure 3c) over the substrate, in line with our hypothesis and previous findings. As for aromatic-rich mPilA strains, pili-abundant regions were observed with clear conductivity increases over that of the background substratum (Figure 2, T61Y; T61W). *CN* T61Y and T61W displayed contact current increases of 0.29 (SD ± 0.08) and 0.36 pA (SD ± 0.06) respectively (Figure 2b), which was 200% greater than the current displayed by the native PilA monomer. This observation was in accordance with pre-existing literature, where PilA modifications to increase tryptophan content have resulted in conductivity increases of 140 – 500% ^5,21^. Filaments with 4.5-5.7nm cross-sectional height and several microns in length were observed in all MPilA strains and control variants (Figure 2c, WT: 5.4nm (SD ± 0.4), Δ*pilA*: 5.3nm (SD ± 0.41), T80: 4.8nm (SD ± 0.55), T61: 4.5nm (SD ± 0.8), T61Y: 5.7nm (SD ± 0.8), WT: 5.6nm (SD ± 0.8)).Unlike observations conducted *in vitro* ^5^, we did not observe a significant (p < 0.05) change in pilus diameter between native and modified filaments, based on one-way ANOVA analysis of control pilus (WT) compared to experimental (Δ*pilA, T80, T61, T61Y, T61W)*.

**Figure 3:**
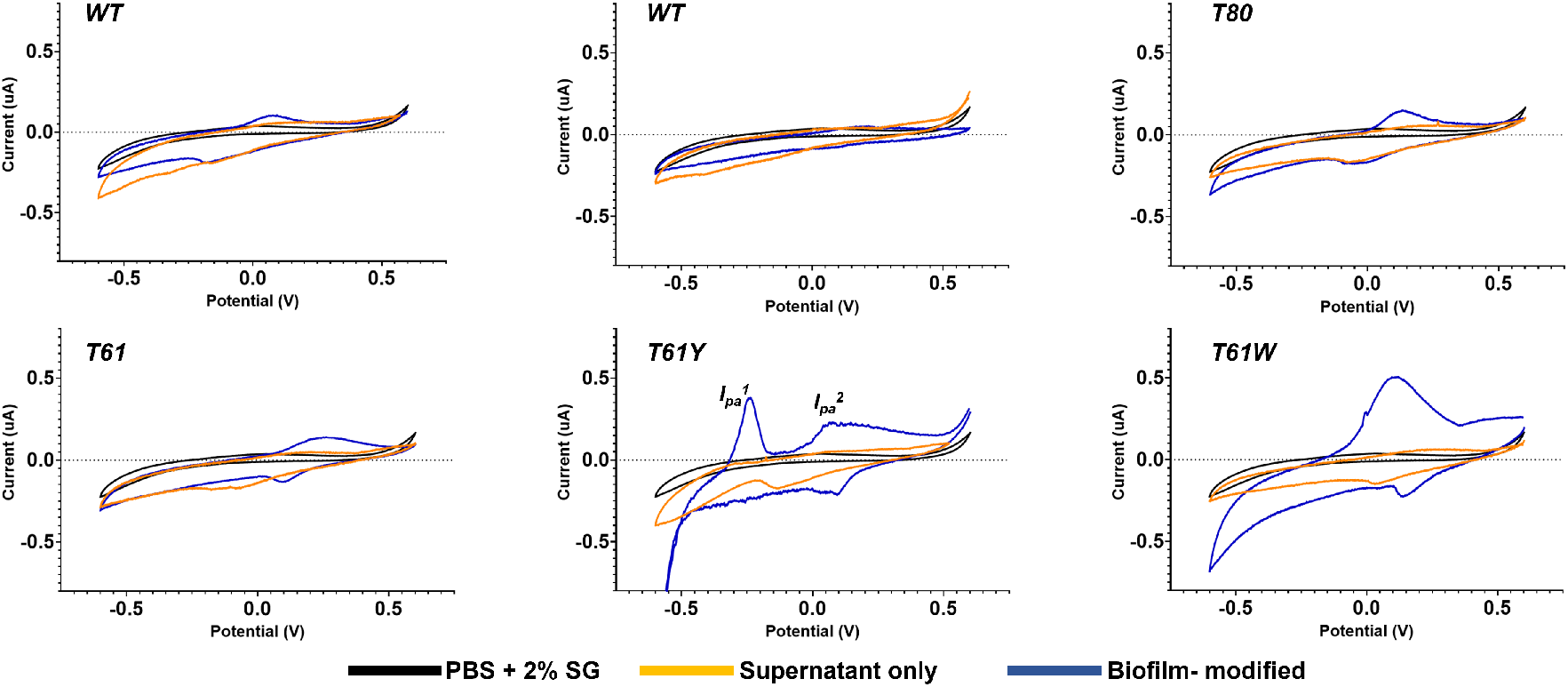
Typical Cyclic Voltammograms of screen-printed carbon electrodes coated with mPilA bacterial biofilms, with the strain labelled at the top-left of each graph. The black trace is the bare medium only (PBS at pH 6.9 + 2% sodium gluconate), the orange trace is cellular supernatant, and the blue trace is a bacterial biofilm-modified electrode. All scans were initiated by performing 50 scans at 100mVs to equalise the electrode surface at 1mVs, followed by two scans at 1mVs with the second scan displayed above. All scans performed in triplicate (n= 3) on separately prepared samples, with the mean peak size and positioning displayed in SI Table 3.

### Cyclic Voltammetry characterisation of the cellular redox behaviour

To investigate if aromatic modification of pilus monomer sequence affects cellular redox behaviour, cyclic voltammetry of biofilm-coated, screen-printed carbon electrodes was performed (Figure 3). For strains without aromatic or structural modifications, oxidative peaks of 0.028µA (SD ± 0.029) and 0.023µA (SD ± 0.01) at, respectively, 0.017V (SD ± 0.03) and 0.2V (SD ± 0.0) were observed for *CN* H16 and *CN* Δ*pilA*. MPilA strains with aromatic modifications, however, displayed varying redox behaviour. In strain T61Y, an additional oxidative peak of 0.23µA (SD ± 0.157) at -0.22v (SD ± 0.029) was present, along with one of 0.15µA (SD ± 0.05) at a similar potential to that of control strains (0.2 V ± 0.1 SD). The strain with increased tryptophan content, *CN T61W* displayed an increased peak oxidative current 9 times larger than that of the WT strain (0.26µA (SD ± 0.11), with a slight shift in peak potential (0.1 V (SD ± 0.1). These values are summarised in SI Table S3.

To examine the origin of the current increases observed with engineered strains, duplicate cyclic voltammetry experiments were performed using only the cellular suspension medium, with cells removed via centrifugation. The increased oxidative peak current detected in aromatic-modified strains was not present in bare medium samples, suggesting that the electrochemistry is not from a solution-based species. This observation indicated that the increased oxidative currents were attributable to cellular-based mechanisms facilitating electrochemical interactions between cells and the electrode, rather than any redox-active soluble mediator secreted into the medium. These results suggest that increased oxidative peak currents may be attributable to the expression of pili with increased aromatic content.

## Discussion

This work is the first example of functional type-IV pili protein nanowires produced under aerobic conditions using a *CN* chassis. Protein nanowires are considered a highly efficient form of long-range direct extracellular electron transport, allowing certain microorganisms to electrochemically connect with the extracellular environment over considerable distances without the use of redox-active secreted mediators. Significantly, GS has achieved the highest current density generation thus far in MFC reactors ^40^ and conductive nanowires are proposed to have a major role in their external electron transfer (EET) capabilities. However, exploiting their industrial useability is complicated by the challenging growth requirements of native hosts, and the debate over their molecular identity is one of the most contentious areas of microbiology.

This work offers two significant contributions. Firstly, we confirm the conductivity of microbial pilin proteins can be engineered via a combination of aromatic content and monomer secondary structure modification. The 200% conductivity increases observed in aromatic-modified PilA strains compared to WT are in line with other researchers’ findings. Furthermore, we demonstrate that the expression of conductive type-IV pili nanowires affects the redox behaviour of recombinant *C. necator* strains, allowing bacterial cells to electrochemically interact with the environment with over 9-fold increases in oxidative peak current compared to the wild-type. Together, these findings provide compelling support for the hypothesis that conductive microbial nanowires are comprised of PilA proteins.

Importantly, the use of *CN* as a chassis for conductive nanowire expression presents further evidence which is hard to reconcile with the suggestion that polymerised cytochrome nanowires were originally misidentified as pili ^15^. A homology study conducted with a protein BLAST alignment against the *CN* H16 non-redundant protein database for the *GS* cytochromes and proteins putatively involved in EET (OmcZ, OmcS, OmcT, OmcE and OmcF), indicated no homology nor sequence similarity within the *CN* proteome (SI, Table 4). Therefore, if conductive nanowires were indeed comprised of stacked OmcS subunits ^10,12,15^, the detection of conductive filaments in our *CN* strains would have been highly unlikely, considering the absence of both proteins with similar binding sites for polymerisation and functional homologs in *CN* ^41,42^.

Secondly, we present the first high-resolution Images which map the current of conductive surface-expressed protein nanowires emanating from intact bacterial cells. In contrast to previous C-AFM observations of pili, including PeakForce TUNA analysis conducted by the sampling of sheared or precipitated surface-expressed filaments, the limited processing and chemical treatment described in this work allows the visualisation and conductivity analysis of pili in conditions close to their native environment. Based on this analysis, we demonstrate that pili conductivity is not attributable to artefacts caused by chemical treatment.

The expression of conductive nanowires in *CN* is a significant development for the intensification of industrial-scale processes using electroactive microbial chassis, with the potential to provide solutions to a range of environmental issues. Type-IV pili are commonly used as adhesive structures by microbial species ^43,44^, expressed by a range of pathogenic organisms to adhere to and colonise host cells ^45^ which complicates their use as an industrially feasible electron transport mechanism. In contrast, *CN* is non-pathogenic and has already been used as a chassis in several pilot-scale industrial projects. These pilot plants aim to produce thousands of metric tons per annum of the degradable bioplastic PHB ^46^, which can then be processed into valuable fuels and chemical feedstocks ^47^, reducing society’s reliance on environmentally damaging fossil fuels. However, current large-scale PHB production systems use agricultural sugars as feedstocks ^48^. Engineering an extracellular electron transfer mechanism into *CN* offers a novel path to CO_2_ remediation and assimilation into useful bioplastics and chemical feedstocks via renewable energy sources. While indirect electron transfer was achieved previously in *CN*, with the addition of redox-active mediators ^49^ and hydrogen production via water electrolysis ^50^, these processes involve additional reaction stages or materials, adding cost and complexity to system design. In this work, we offer the first example of a purely biological, direct electron transfer mechanism achieved in *CN* based on type-IV pili nanowires. These findings offer new opportunities for alleviating global fossil fuel reliance *via* the production of sustainable biofuels based on the reduction of atmospheric carbon.

## Materials and Methods

### Generation of the *Cupriavidus necator* H16 Δ*pilA* chassis

*Cupriavidus necator* H16 was obtained from the University of Nottingham’s Synthetic Biology Research Centre culture collection. *CN’s* native *pilA* sequence was identified (Q0KE72, genomic location 577442:577936), and 1KB fragments of the locations immediately upstream and downstream were amplified using the primers HIFI:PilAKO:UpHA:FWD/ REV and PilAKO:DownHA:FWD/ REV (SI Table S1), respectively. These fragments were then assembled onto the pMTL70641 suicide plasmid, previously described ^34,51^ containing the tetracycline resistance cassette and *sacB3* gene via a three-fragment HiFi reaction (New England Biolabs). *Escherichia coli* S17-1 was transformed with the pMTL70641:*ΔpilA* construct and conjugated with *C. necator* H16 for six hours at 30°C. Successful conjugants were purified by growing on selective sodium gluconate minimal medium ^52^ agar plates containing 15ug/ml gentamicin and tetracycline (SGMMGT) overnight at 30°C. Purification was repeated and after a second overnight incubation, colonies were re-plated on replicate low-salt LB plates containing 15% sucrose, and SGMMGT plates. 200 colonies were plated and incubated at 30°C for 48 hours. Colonies exhibiting growth on the 15% sucrose plate, but not on the selective plate were re-streaked on a SGMM agar plate containing 15ug/ml gentamicin. Diagnostic colony PCR reactions were performed, to identify if counter-selection had occurred successfully and to determine if colonies contained the deletion region. Once the colonies had passed screening, they were re-purified on selective minimal medium agar plates and stored in glycerol at -80°C. A final verification stage was performed via Next-generation sequencing of the whole genome (MicrobesNG), which confirmed the absence of the pilA coding region.

### Modified PilA (mPilA) sequence design

*CN’s* native PilA monomer sequence (Q0KE72) was adapted to produce a range of synthetic fragments, potentially encoding for conductive PilA monomers. The modified PilA sequences were: T80, truncated to 80 amino acid residues in length leaving a portion of C-terminal β-sheet; T61, further truncated to a 61 amino acid length, leaving only α-helix; T61-Y, T61 with the native amino acids at positions 32, 51, 51 (arginine; alanine; asparagine) replaced for aromatic (tyrosine; phenylalanine; tyrosine) respectively; T61-W; an adaptation of T61-Y with the same three amino acids from the previous fragment replaced with the aromatic tryptophan. All synthetic PilA fragments were preceded by *CN’s* native MQRVQQLKKLGRRVQKG signal sequence, to allow recognition of the translated protein, transportation to type 4 construction apparatus and secretion through the membrane. All fragments included the 6*HIS tag directly at the N-terminus, followed by the SPC protein of *GS*. Fragments were synthesised by GeneWiz (NJ, USA).

### Plasmid assembly

The plasmid pMTL71301, reported previously ^53^ was sourced from within the lab group and digested via the restriction enzymes *BgI*II;, *Hin*dIII (New England Biolabs). *mpilA* fragments were amplified with overhanging regions of 25 base pairs to allow integration within the assembled construct, using forward and reverse primers HiFi:71301:MPilA:ALL:FWD and HiFi:71301:MPilA:ALL:REV (SI Table S1) respectively, and purified via Monarch gel extraction kit (New England Biolabs). The *araBAD* inducible promoter ^54^ was ligated to the 5’ end of the plasmid and mPilA fragments were located immediately downstream. All ligations were performed via HiFi master mix two-fragment assemblies (New England Biolabs). The assembled pMTL71301: *araBAD:mpilA* constructs were verified *via* Sanger sequencing (Eurofins).

### Transformation and verification

*C. necator* H16*:ΔpilA* electrocompetent cells were produced by incubating at 30°C overnight in Super Optimal Broth medium (Sigma-Aldrich), harvested by centrifugation and rinsed twice in 1ml of 1mM MgSO_4_. Electrocompetent cells were resuspended in 50μL of 1mM MgSO_4_ and mixed with approx. 250ng of plasmid DNA. Electroporation was performed in a pre-chilled 2mm gap cuvette (Scientific Laboratory Supplies) at 2.5kV, 200Ω and 25µF (Bio-Rad). 0.95ml of Super Optimal Broth with Catabolites (New England Biolabs) was added immediately after electroporation and incubated at 30°C with rapid shaking for 60 minutes. 100μL of each cell suspension was plated on nutrient-limiting agar plates containing sodium gluconate as a carbon source (SGMM), supplemented with gentamicin (to inhibit *E. coli*) and tetracycline to eliminate the growth of wild-type *C. necator* cells. Sodium Gluconate minimal medium with gentamicin and tetracycline (SGMMGT) plates were incubated at 30°C until colonies were visible (approx. 48-72 hours). Presumptive screening of 10 colonies per mPilA variant was performed via colony-PCR using GoTaq® Green Master Mix (Promega) and forward and reverse primers Diag:71301:MPilA:All:FWD and Diag:71301:MPilA:All:REV (SI Table S1), respectively. Each colony was re-streaked on the same selective SGMMGTT agar plates, and the loop was used to inoculate the colony PCR tube. Plates were grown at 30°C for 48-72 hours. Overnight SGMMGT broth cultures of colonies containing the complete 1.6kB mPilA construct were grown at 30°C for approx. 16 hours and plasmid DNA was extracted using the QIAprep Spin Miniprep Kit (Qiagen). Final verification was performed via Sanger sequencing (Eurofins), using the forward and reverse Diag:71301:MPilA:All:FWD and Diag:71301:MPilA:All:REV (SI Table S1) respectively, to verify the success of electroporation and integration of the plasmid containing the correct mPilA fragment. At this stage, a complete set of *C. necator* H16:*ΔpilA:mpilA* variants had successfully passed both presumptive colony PCR diagnostics and sanger sequencing verification stages. Glycerol stocks of all successful variants were inoculated and stored at -80°C until use.

### Western blot immunodetection of PilA expression

PilA monomers have a molecular weight of 8-15 kDa ^6 21^. To resolve proteins of this low molecular weight, 10-20% Tricine NuPage gel system was used (Thermo-Fischer).

*C. necator* H16*:ΔpilA: mpilA* strains (T80; T61; T61Y; T61W), along with a *C. necator* H16*:ΔpilA* strain containing an empty pMTL71301: mPilA expression plasmid without the mPilA cargo were grown in SGMMGT broth ^52^ overnight at 30°C. Following this, pili and surface filaments were physically sheared for SDS-PAGE-Western blotting as in the protocol previously described ^55^. 50µL of culture broth was spread-plated on SGMMGT agar plates containing 5mM Arabinose to form a dense bacterial lawn under pilus-inducing conditions and incubated at 30°C for 24 hours. A large loop of bacterial cells was resuspended in 1ml of sterile PBS (Phosphate Buffered Saline) and normalised to an OD600 of 1. The cell suspension was vortexed at high speed for two minutes and passed through a 26-gauge needle six times, to lyse the cells and detach surface filaments as in a protocol previously published ^37^. The lysate was centrifuged for 15 minutes at 16000rcf, and the supernatant was decanted to produce sheared fragment (SF). This was centrifuged for a further 10 minutes at 16000rcf to remove any residual cellular debris and precipitated in 10% Trichloroacetic acid on ice for 60 minutes. The precipitate was centrifuged at 16000rcf for 60 minutes, the supernatant discarded, and the pellet washed with ice-cold acetone twice. After drying in a heat block at 70°C for 10 minutes, the pellet was resuspended in 70ul of Novex® sample buffer (Invitrogen). Nupage reducing agent (Thermo-Fisher) was added to all samples at 1:10 and heated to 70°C for 10 minutes to denature proteins. After heating, samples were placed in ice for 5 minutes and loaded into a 10-20% Tricine gel cassette along with Spectra low-range protein ladder and the BenchMark™ His-tagged Protein Standard (Thermo-Fisher). SDS-PAGE was performed at 120V for approx. 120 minutes.

The separated proteins present on the SDS-PAGE gel were transferred to a nitrocellulose membrane using the BioRad Transblot instrument set at the “Low molecular weight” programme (5V, 5 minutes). The protein-loaded membrane was immediately submerged in Tris-Buffered Saline + 1% Tween-20 (TBS-T) blocking solution containing 5% Bovine Serum Albumen (BSA) to block unspecific antibody binding to the non-target proteins and incubated at room temperature for 60 minutes.

After this time, anti-HIS tag primary antibody (Thermo-Fischer) was added at the manufacturer’s recommended ratio of 1:5000 and incubated at 4°C overnight. The primary antibody solution was decanted, and the membrane was rinsed three times in TBS-T for 15 minutes with gentle agitation to remove residual primary antibodies. The membrane was submerged in secondary antibody solution (TBS-T + 5% BSA + 1:3000 Anti-Mouse IgG (Fab specific)-Peroxidase antibody (Sigma-Aldrich) and incubated for two hours at room temperature. The membrane was rinsed three times for 30 minutes in TBS-T, and protein bands were visualised via coating the membrane in tetramethylbenzidine (Sigma-Aldrich).

### PeakForce TUNA Atomic Force Microscopy

PeakForce TUNA AFM was used to image and perform conductivity analysis of pili. Glass coverslips were used as substrates for bacterial growth and AFM imaging. Coverslips were cleaned by bath sonication, using alternate washes of ethanol: HCL (v/v 70:1) ^56^ and sterile RO water for 10 minutes. The process was repeated for a total of two wash cycles, and coverslips were dried with a stream of nitrogen in a fume cupboard. *CN* H16:*ΔpilA: mpilA* variants were revived from -80°C freezer stocks by plating on SGMMGT plates containing 1.5% bacterial agar #1 (Oxoid) and incubated for 24-48 hours at 30°C. SGMMGT broth was inoculated with a plated colony and incubated overnight at 30°C, with 20μl of the resulting culture used to inoculate a 6-well plate containing 3ml of SGMMGT broth and a cleaned glass coverslip. The plate was incubated at 30°C for 24 hours, with pili expression induced via the addition of 5mM Arabinose after 6 hours of growth. After 24 hours, the culture broth was decanted, coverslips rinsed gently using sterile RO water to remove planktonic cells and dried via a stream of nitrogen in a fume cupboard. Imaging was performed on a Dimension Icon AFM (Bruker) using PeakForce Tapping^TM 57^ with a PeakForce TUNA module. Platinum-coated conductive AFM cantilever tips with a resonant frequency of 150 kHz (NuNano) were used to image bacterial cells and surface filaments. To find regions of interest a low image resolution of 200 samples/ line was used, then increased to 500 when a satisfactory field of view was reached. Instrument parameters used included a scan rate of 1.5 Hz, peak force amplitude of 200nm, current bias and sensitivity of 10 V; 20 pA/V respectively.

### Cyclic Voltammetry characterisation of the cellular redox behaviour

*C. necator* strains were revived from -80^°^C freezer stocks by plating on SGMMGT agar plates and incubating for 24-48 hours at 30°C. Overnight broth cultures were inoculated with single colonies and incubated at 30°C for 16 hours, with MPilA expression induced via the addition of 5uM arabinose after 6 hours of growth. The pellet was washed with PBS (pH adjusted to 6.9) a total of two times and resuspended to an OD_600_ of 3.5 in PBS, pH 6.9. A Zensor TE-100 3-electrode carbon screen-printed carbon electrode (SPCE) was rinsed with sterile water, placed in a culture tube along with the standardised culture, sealed with parafilm and left in an anaerobic cabinet (Don Whitley Scientific) overnight to degas the culture of dissolved O_2_. After 16-24 hours the biofilm-modified electrode was removed from the culture tube, gently rinsed with sterile water to remove planktonic cells and connected via crocodile clips to the potentiostat (Metrohm). To equalise the electrode surface, 50 scans at 100mV s^-1^ were performed from 0.6 V to -0.6 V, followed by two scans at 1mVs to capture the redox behaviour at the electrode-biofilm interface. To determine if any redox peaks observed were attributable to soluble redox-active compounds secreted into the culture media, duplicate experiments were performed including an additional centrifugation step (10,000 rcf, 10 minutes) after cell density standardisation to remove all cellular material with the electrode submerged in supernatant only. A total of three repeats were performed for each bacterial strain, from distinct colonies. Data analysis was performed via DropSens software (Metrohm)

## Data availability

All data comprising this manuscript is available https://rdmc.nottingham.ac.uk/

## Acknowledgements

This work was supported by the Engineering and Physical Sciences Research Council (grant numbers EP/R004072/1, EP/S017739/1), We thank the Nottingham-Rothamstead Doctoral Training Partnership for providing a studentship to B.M. and F.C. (BB/M008770/1, Project reference 2275606 and 1943801). In addition it was funded by the Biotechnology and Biological Sciences Research Council (grant number BB/L013940/1 (BBSRC); and the Engineering and Physical Sciences Research Council (EPSRC) under the same grant number.

## Contributions

FJR., PJH, and KK conceived the ideas in the project. BM., FC., SA., PJH., KK., FJR. designed the experiments. BM performed all experiments and processed the data, BM., SA., PJH., and KK., FJR. analysed the data. BM., FC, SA, PJH., and KK., FJR wrote the manuscript and FC, SA, PJH., and KK. edited.

