## Supporting information for "Engineering Nanowires in Bacteria to Elucidate Electron Transport Structural-Functional Relationships"

Ben Myers[1,3], Francesco Catrambone[2], Stephanie Allen[3], Phil J Hill[4], Katalin Kovacs[1,3*], Frankie J Rawson[1*]

[1] Bioelectronics Laboratory, Regenerative Medicine and Cellular Therapies, School of Pharmacy, Biodiscovery Institute, The University of Nottingham, Nottingham, NG7 2RD, UK

[2] BBSRC/EPSRC Synthetic Biology Research Centre, School of Life Sciences, Biodiscovery Institute, University of Nottingham, NG7 2RD, United Kingdom

[3] Molecular Therapeutics and Formulation Division, School of Pharmacy, Boots Science Building, University Park, The University of Nottingham, Nottingham, NG7 2RD, UK

[4] Division of School of Microbiology, Brewing and Biotechnology, School of Biosciences, Sutton Bonington Campus, University of Nottingham, LE12 5RD, United Kingdom

### Supplementary Methods:

**Table S1:** Table of plasmids used in this study

| Plasmid | Function | Source |
|---|---|---|
| pMTL70641:SacB3 | Suicide plasmid host vector | Muhammad Ehsaan [1] |
| pMTL70641:SacB3:PilAKO | Suicide plasmid (PilA deletion) | This study |
| pMTL71301: araBAD | MPilA expression host vector | Muhammad Ehsaan [1] |
| pMTL71301:araBAD:mPilA:Empty | Modified PilA negative control (no MpilA cargo) | This study |
| pMTL71301:araBAD:mPilA:T80 | Modified PilA expression (strain T80) | This study |
| pMTL71301:araBAD:mPilA:T61 | Modified PilA expression (strain T61) | This study |
| pMTL71301:araBAD:mPilA:T61Y | Modified PilA expression (strain T61Y) | This study |
| pMTL71301:araBAD:mPilA:T61W | Modified PilA expression (strain T61W) | This study |

**Table S2:** Table of primers used in this study

| Primer | Sequence (5"-3") | Function |
|---|---|---|
| HIFI:PilAKO:UpHA:FWD | TTCGAGCTCGGTACCCGGGGATCCTCTTC CACCACATTCTGGATTG | Amplification/ HIFI Assembly |
| HIFI:PilAKO:UpHA:REV | GGCATTCCGTACGATTTGACCCCTCGAAG AG | Amplification/ HIFI Assembly |
| PilAKO:DownHA:FWD | GAGGGGTCAAATCGTACGGAATGCCTGA GAAAAAGCGCCTC | Amplification/ HIFI Assembly |
| PilAKO:DownHA:REV | TGCCAAGCTTGCATGTCTGCAGGCCTGCG CCGCGTGCGGACAG | Amplification/ HIFI Assembly |
| Diag:PilAKO:Ext:FWD | TTCCAGCGAGCGCTGGAATGGCGAA | PCR Diagnostics |
| Diag:PilAKO:Ext:REV | GACTTGCTGCAACTGCTGGAATTGCGC | PCR Diagnostics |
| Seq:PilAKO:Up:FWD | CTTCCACCACATTCTGGATTG | Sequencing |
| Seq:PilAKO:Int:FWD | ATGCAACGGGTACAACAACTG | Sequencing |
| Seq:PilAKO:Down:REV | GATAGCGAAGAGCGCAAGCA | Sequencing |
| HiFi:71301:MPilA:ALL:FWD | CGACGTCACGCGTCCATGGAACCGCGGCC GCTGTCAAA | MPilA Insert HiFi Assembly |
| HiFi:71301:MPilA:ALL:REV | CGATGACGACAAGTAATAAGCGCTAGCAT TGGCACTGGCCGTCGTTTTA | MPilA Insert HiFi Assembly |
| Diag:71301:MPilA:All:FWD | CGCTTCAGCCATACTTTTCATAC | PCR Diagnostics/ Sequencing |
| Diag:71301:MPilA:All:REV | GCTCTTGGATGGAGGAAATGAC | PCR Diagnostics/ Sequencing |

**Table S3:** Mean peak size and potential of biofilm- modified electrodes, detected via cyclic voltammetry exemplified in figure 5 of the main manuscript.

| | **Oxidative Peak Current (µA)** $\mathbf{I_{pa}}$ | **Oxidative Peak Potential (V)** $\mathbf{E_{pa}}$ | **Reductive Peak Current (µA)** $\mathbf{I_{pc}}$ | **Reductive Peak Potential (V)** $\mathbf{E_{pc}}$ |
|---|---|---|---|---|
| **WT** | 0.028 (+/- 0.029) | 0.017 (+/- 0.03) | -0.025 (+/- 0.035) | 0.075 (+/-0.23) |
| **ΔpilA** | 0.023 (+/- 0.01) | 0.2 (+/- 0.0) | -0.015(+/- 0.03) | 0.05 (+/- 0.05) |
| **T80** | 0.062 (+/- 0.041) | 0.17 (+/- 0.06) | -0.03 (+/- 0.02) | -0.05 (+/- 0.08) |
| **T61** | 0.04 (+/- 0.03) | 0.23 (+/- 0.058) | -0.039 (+/- 0.02) | 0.03 (+/- 0.058) |
| **T61Y** | 1: 0.23 (+/- 0.157)<br>2: 0.15 (+/- 0.05) | 1: -0.22 (+/- 0.029)<br>2: 0.2 (+/- 0.1) | -0.05 (+/- 0.038) | 0.2 (+/- 0.09) |
| **T61W** | 0.26 (+/- 0.11) | 0.1 (+/- 0.1) | -0.11 (+/- 0.1) | 0.02 (+/- 0.11) |

**Table S4:** Cytochrome BLASTp sequence alignment within all non-redundant GenBank CDS translations + PDB + SwissProt + PIR + PRF excluding environmental samples from WGS projects databases. Queried microorganisms: *G. sulfurreducens* KN400 (taxid:663917), Geobacter sulfurreducens (strain ATCC 51573) and *C. necator* H16 (taxid:381666).

| Description | Scientific name / Taxid | Max score | Query cover | E value | Identity | Acc. Length | Gene ID: |
|---|---|---|---|---|---|---|---|
| OmcS | Geobacter sulfurreducens (strain ATCC 51573) | No significant similarity found | n/a | n/a | n/a | 432 | Q74A86 |
| OmcB | Geobacter sulfurreducens (strain DL-1 / KN400) | No significant similarity found | n/a | n/a | n/a | 744 | D7ALQ0 |
| OmcZ | Geobacter sulfurreducens (strain ATCC 51573) | No significant similarity found | n/a | n/a | n/a | 473 | Q74BG5 |
| OmcT | Geobacter sulfurreducens (strain ATCC 51573) | No significant similarity found | n/a | n/a | n/a | 430 | Q74A87 |
| OmcE | Geobacter sulfurreducens (strain ATCC 51573) | No significant similarity found | n/a | n/a | n/a | 232 | Q74FJ0 |
| OmcF | Geobacter sulfurreducens (strain ATCC 51573) | No significant similarity found | n/a | n/a | n/a | 104 | Q74AE4 |
| PgcA | Geobacter sulfurreducens (strain ATCC 51573) | 32.7 | 6% | 0.033 | 38.71 | 511 | Q74CB3 |
| Cytochrome c family protein | Cupriavidus necator H16 | 32.7 | 6% | 0.033 | 38.71 | 125 | WP_011617786.1 |

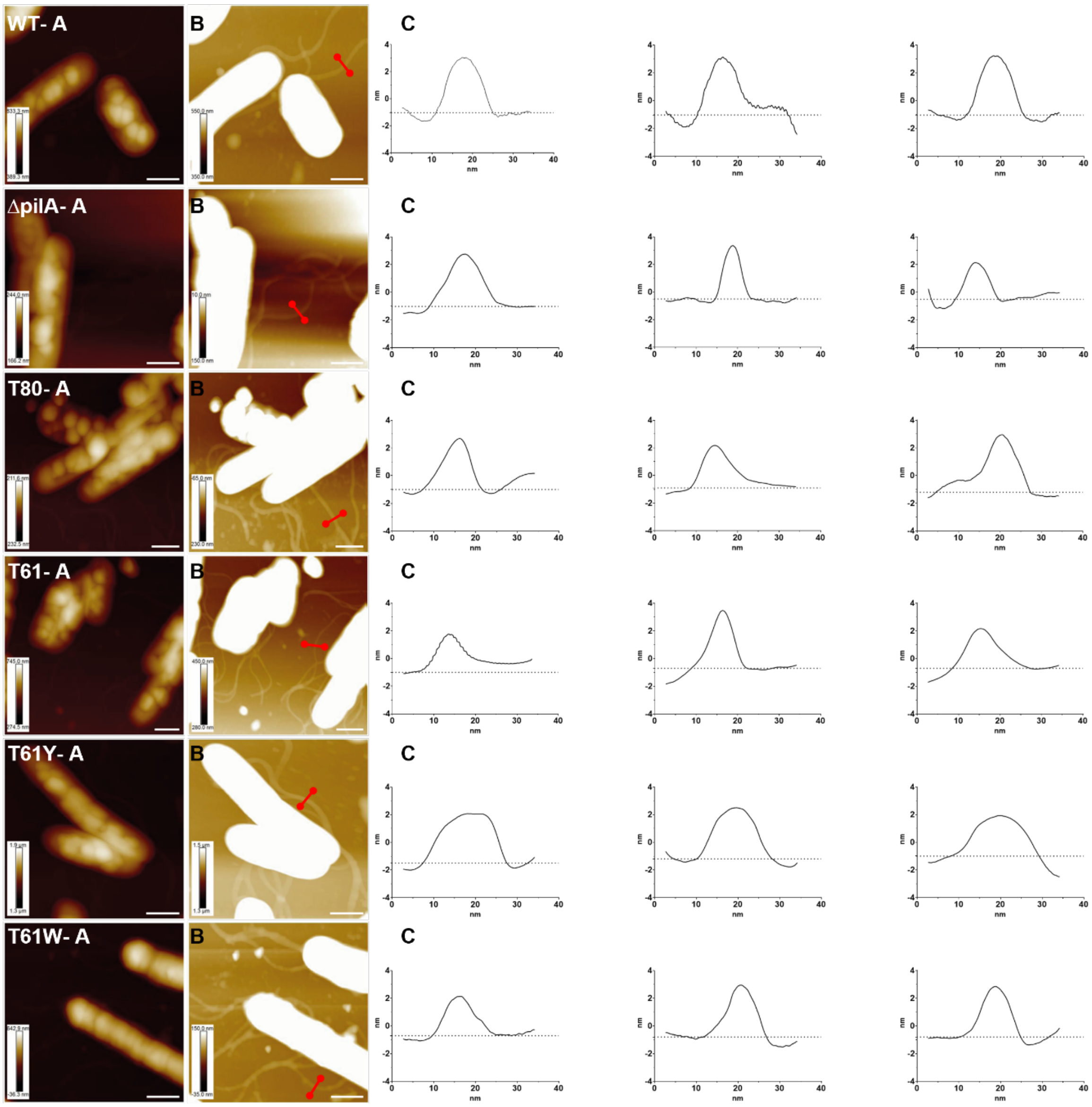


**Figure S1: Analysis of AFM topographical data to determine pili morphology.** A: Raw topographical (height) data map of *CN* strains WT- T61W, with z- scale automatically determined via Nanoscope analysis software. Due to large height differential between bacterial cells and filaments, nanoscale features are difficult to observe in the raw z- scale and so manual narrowing was required. B: Topographical data map with z- scale manually decreased to bring features within range to allow visualisation. Z- scale shown as insert in bottom- left, and white bar in bottom right representing a lateral scale of 1 µm in both A and B. Red cross sectional bar indicating representative locations for height profile gradient analysis in B. C: Line profile analysis of pili- cross sectional height, generated via Gwyddion. Each graph displays the mean of three measurements, taken across an individual filament. Three filaments were analysed for each strain, making a total n of 9 measurements. The baseline used to calculate filament height is displayed as the dotted line within each graph. All variants had a mean filament morphology of 4.5-5.7nm (WT: 5.4nm SD ± 0.4, Δ*pilA*: 5.3nm SD ± 0.41, T80: 4.8nm SD ± 0.55, T61: 4.5nm SD ± 0.8, T61Y: 5.7nm SD ± 0.8, WT: 5.6nm SD ± 0.8. Differences in recombinant pili morphology were not significant ($p>0.05$), based on one- way ANOVA analysis of control pilus (WT) compared to experimental (*ΔpilA, T80, T61, T61Y, T61W)*. Data analysis performed via Graphpad prism.

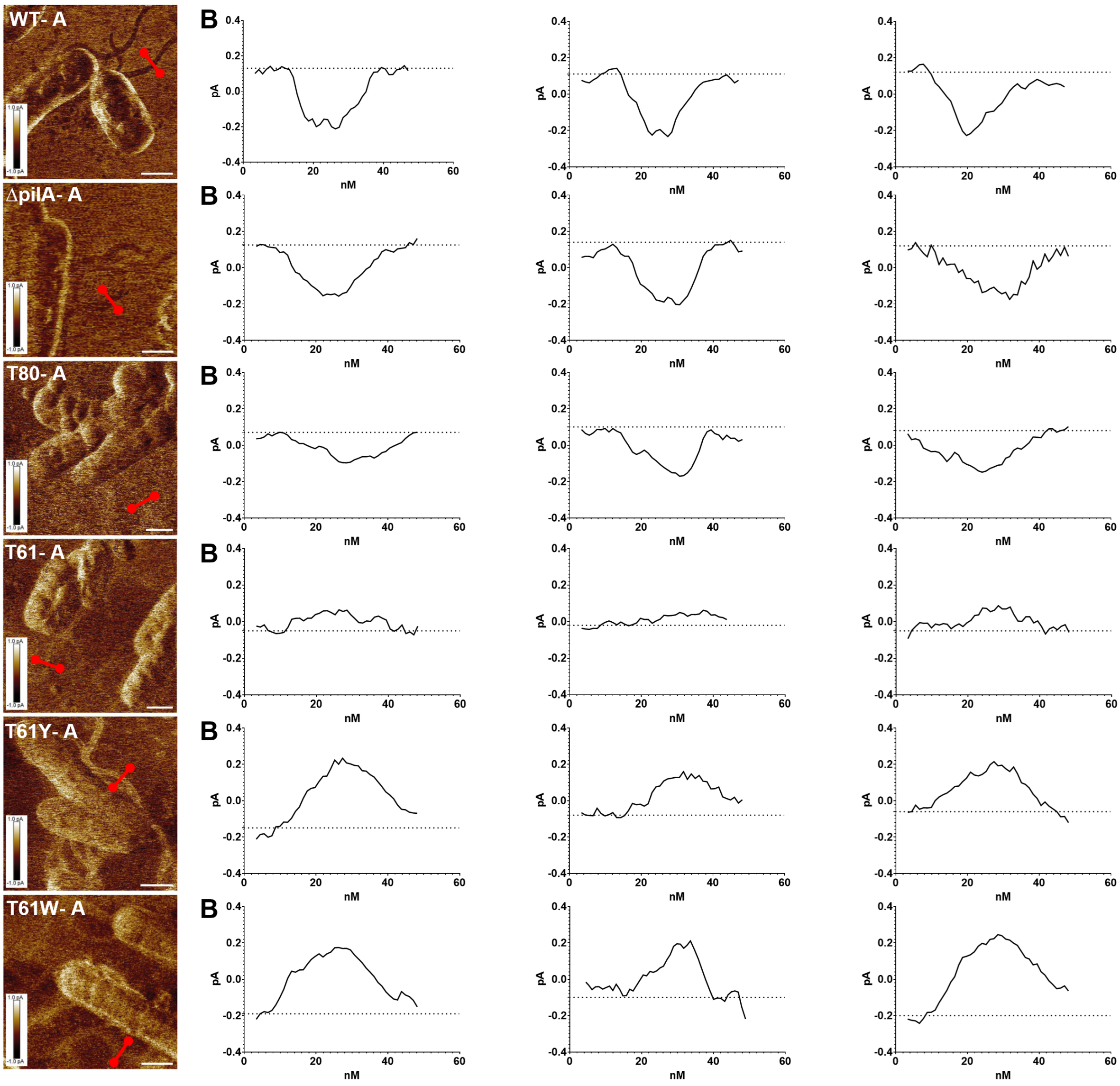


**Figure S2. AFM Contact Current data analysis to determine current profile of pili.** A: Contact current data map of *CN* strains WT- T61W. Z- scale shown as insert in bottom- left, aligned across all data maps to a scale of +1pA to -1pA and white bar in bottom right representing a lateral scale of 1 µm. Red cross sectional bar indicating representative locations for contact current profile gradient analysis. B: Contact current line profile gradients for *CN* strains WT- T61W respectively. Each graph displays the mean of three measurements taken across an individual filament. Three filaments were analysed for each strain, making a total n of 9 measurements. Line profile analysis of pili- cross sectional contact current generated via Gwyddion[2]. The baseline used to calculate filament conductivity is displayed as the dotted line within each graph. WT:-0.33pA SD ± 0.01; ΔpilA:-0.31pA SD ± 0.03; ΔpilA:mPilAT80: 0.21pA SD ± 0.05; ΔpilA:mPilAT61: 0.09pA SD ± 0.03; ΔpilA:mPilAT61Y: 0.29pA SD ± 0.08; ΔpilA:mPilAT61W: 0.36pA SD ± 0.06. Data analysis performed via GraphPad Prism.
